# Parallel Evolution in the Emergence of Highly Pathogenic Avian Influenza A Viruses

**DOI:** 10.1101/370015

**Authors:** Marina Escalera-Zamudio, Michael Golden, Bernardo Gutiérrez, Julien Thézé, Jeremy Russell Keown, Loic Carrique, Thomas A. Bowden, Oliver G. Pybus

**Author notes:** MEZ and MG contributed equally to this paper. other email addresses.

## Abstract

Parallel molecular evolution and adaptation are important phenomena commonly observed in viruses. Here we exploit parallel molecular evolution to understand virulence evolution in avian influenza viruses (AIV). Highly-pathogenic AIVs independently evolve from low-pathogenic ancestors via acquisition of a polybasic cleavage sites (pCS). Why some AIV lineages but not others evolve in this way is unknown. We hypothesise that the parallel emergence of highly-pathogenic AIV may be facilitated by permissive or compensatory mutations occurring across the AIV genome. We combined phylogenetic, statistical and structural approaches to discover parallel mutations in AIV genomes associated with the highly-pathogenic phenotype. Parallel mutations were screened using a new statistical test of mutation-phenotype association and further evaluated in the contexts of positive selection and protein function. The mutational panel we present reveals new links between virulence evolution and other viral traits and raises the possibility of predicting aspects of AIV evolution.

## INTRODUCTION

Avian Influenza A viruses (AIVs) infect bird populations worldwide ^1^. While most AIVs cause only a mild-to-asymptomatic infection, highly-pathogenic (HP) avian influenza, restricted to the H5 and H7 subtypes of AIV, can cause extremely high mortality (up to 100%) in domestic bird populations ^1^. Both low-pathogenic (LP) and HP viruses can cause severe disease in humans following avian-to-human transmission, and there are potential instances of human-to-human transmission ^2,3^. Recently, the emergence of H7N9 AIV in China caused >1500 infections and 615 deaths in people between 2013 and early 2018 ^4^. Low virulence AIVs have evolved into HP strains on multiple occasions, resulting in geographically and temporally distinct HP outbreaks that generate sizeable economic losses to the poultry industry, and pose an increased risk for avian-to-human spread ^5 6^. Therefore, it is important to understand the mechanisms driving the evolution of virulence in AIVs.

The virulence of an infectious disease is, in general, a complex trait ^7 8 9^. However, the AIV HP phenotype is one of the simplest and best characterized molecular markers of virulence, and is therefore an attractive system in which to study virulence evolution. The HP phenotype is conferred by a polybasic cleavage site (pCS) within the virus haemagglutinin (HA) protein. A pCS is the outcome of one or more insertion mutations resulting in multiple basic amino acids, which renders the HA protein a target for broadly-expressed host proteases, facilitating systemic virus spread within avian hosts ^10 11^. The pCS insertion has occurred several times during the evolutionary history of AIVs. Over ten different pCS sequences have been described for HP H5NX viruses since 1961, and over fifteen pCS have been described for HP H7NX viruses since 1963 ^12^. The evolutionary pathways that lead to the independent, parallel evolution of pCSs and the emergence of the HP phenotype in different AIV lineages are not well understood.

Parallel molecular evolution describes repeated evolutionary changes leading to the same phenotype or genotype in independent populations, and can result from adaptation by natural selection to similar selective pressures or ecological niches ^13,14^. RNA viruses exhibit high rates of mutation and replication, large population sizes, consistently varying environments, and strong selective pressures. These factors, combined with small genome sizes, which may restrict the number of mutational pathways leading to a given genotype, may result in parallel molecular evolution being common in RNA viruses ^15^.

We hypothesize that the parallel evolution of the HP genotype from LP ancestors in AIV subtypes H5 and H7 may have a genetic basis, and may be therefore associated with permissive or compensatory secondary mutations occurring elsewhere in the virus genome. Such mutations could arise prior to or immediately after the emergence of a pCS, perhaps to compensate for possible detrimental effects on the stability of the HA protein ^16,17^. Little is known about the genetic predisposition of AIV to acquire a pCS (except for RNA structural studies of the cleavage site region itself ^18^), nor about the role of mutations elsewhere in the virus genome favouring the evolution to a HP phenotype.

Here we use a combination of analytical methods from evolutionary and structural biology to detect parallel mutations across the AIV genome that are associated with the evolution of the HP phenotype (hereafter termed *HP-cluster associated parallel mutations, or HAPMs*). We developed, validated and implemented a new phylogenetic model of binary trait evolution that tests whether genotype-to-phenotype associations observed across multiple lineages are statistically significant. This approach explicitly incorporates the phylogenetic correlation among observations and thus is less susceptible to error than standard statistical tests ^19^. We further investigated if the HAPMs detected had evolved under positive selection, and assessed their possible effects on protein function and/or stability. We identify numerous HAPMs among HP AIV lineages of same viral subtype, many of which are also associated with other viral processes, including antigenic escape, host-response modulation, and replication efficiency. We discuss how the mutational panel presented here can provide a starting point for future molecular studies on the evolution of the HP phenotype.

## METHODS

We developed a new analysis strategy to detect parallel genetic changes associated with the LP-to-HP evolution of AIV (summarized in Figure 1). For convenience, we refer to parallel genetic changes as HAPMs (*HP-cluster associated parallel mutations*). The list of HAPMs identified for each virus subtype was further analysed to identify those most likely to be functionally and selectively relevant using the following criteria: *i)* they exhibit a statistically-supported association with the HP phenotype, *ii*) they show evidence of being evolving under positive selection, *iii)* they are predicted to be stabilizing to protein structure, and *iv)* they have known or predicted functional relevance, derived from structural analysis.

**Figure 1.**
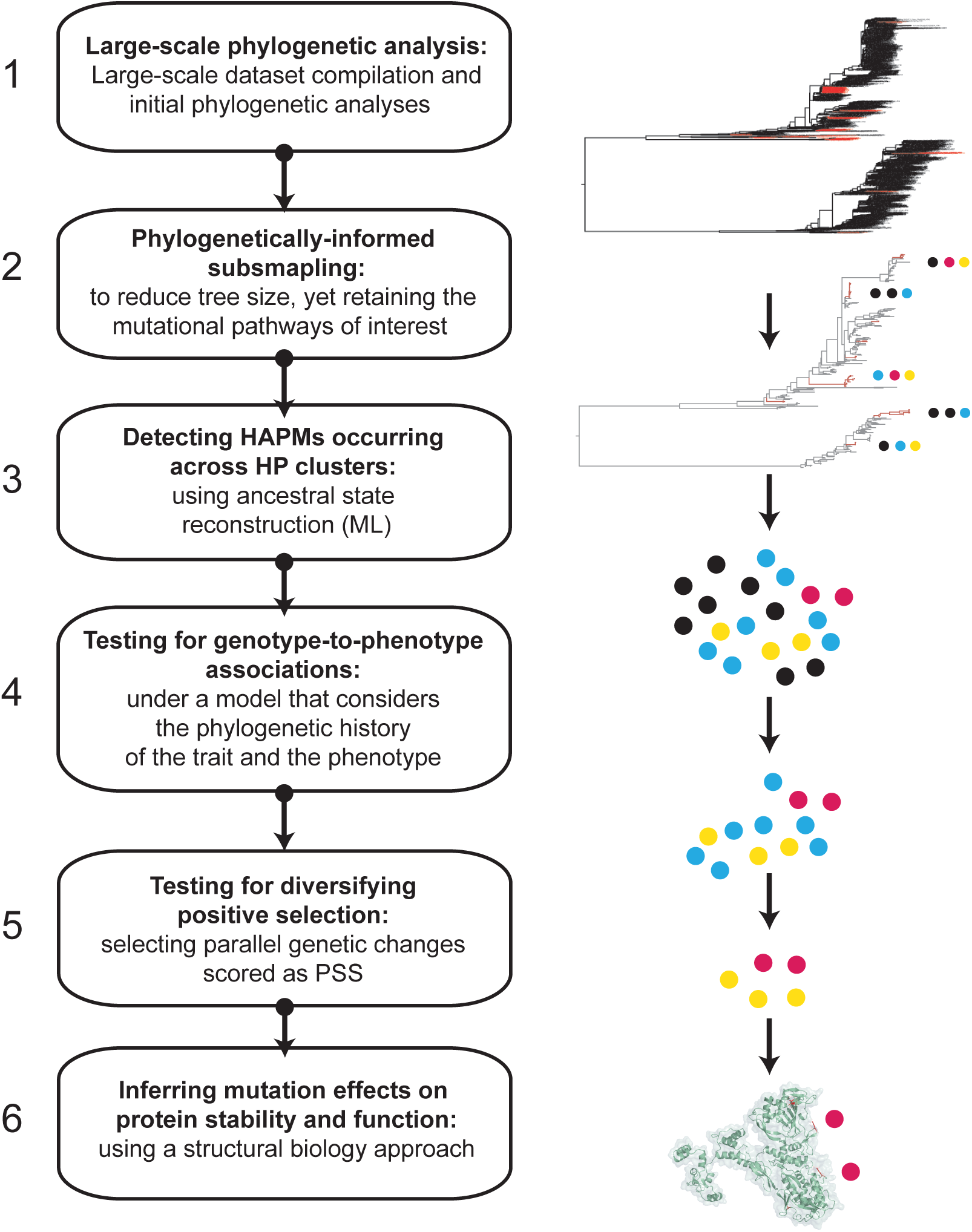
Summary of analytical approach. A summary of our approach to identifying HAPMs (HP-cluster associated parallel mutations) in AIV genomes. 1. Large-scale data set curation and phylogenetic analysis to detect genotypes and phenotypes of interest (HP lineages highlighted in red). 2. Phylogenetically-informed subsampling to reduce alignment size whilst retaining key mutational pathways along internal branches. 3. Detecting HAPMs across multiple HP clusters using ancestral state reconstructions (HAPMs shared among clusters represented by coloured dots) 4. Statistical testing for genotype-to-phenotype associations. 5. Identifying HAPMs evolving under positive selection using different *dN/dS* (ω) estimation methods. 6. Inferring the possible effects of HAPMs on protein stability and function by inferring ddG values, and by using existing structural data to infer interactions.

### 1. Large-scale phylogenetic analysis

All available coding sequences for all genome segments of H7NX and H5NX viruses, from all hosts and geographical regions, were downloaded from the Influenza Virus Database. Sequences were excluded if (i) they were >200nt shorter than the full coding sequence length, (ii) they represented identical sequences, or (iii) they contained >10% ambiguities. Sequences were aligned using MAFFT ^20^ and preliminary large-scale phylogenies for each genome segment were estimated using RAxML under a general time reversible nucleotide substitution model with gamma distributed among-site rate variation (GTR+G). Branch support was assessed using a bootstrap approach with 100 replicates ^21^. HP strains were identified by (i) detecting >1 basic residue insertions within the consensus cleavage site in the HA protein alignments ^12^ and (ii) by confirming HP status with case reports in the literature ^5^. Phylogenetic clusters of HP sequences were identified as (i) monophyletic and well supported (bootstrap value >80) clusters that, (ii) contain >2 HP sequences, (iii) descend from LP ancestors in all genome segments, and (iii) have emerged independently from other HP clusters, and are thus distinct in time and space.

### 2. Phylogenetically-informed subsampling

To render further analyses computationally feasible, we sub-sampled the abovementioned datasets in a phylogenetically-informed manner. Subsampling was undertaken so as to retain the mutational pathways along internal tree branches that lead from LP ancestors to HP clusters, whilst reducing overall alignment sizes. For each genome segment, we retained (i) selected HP clusters, (ii) their immediate LP ancestors in the tree, (iii) up to 250 randomly-selected LP reference sequences, in order to maintain overall tree topology, and (iv) a group of ancestral sequences for rooting purposes. Sub-sampled datasets were re-aligned and screened for recombination using the RDP, GENECOV and BOOTSCAN components of the RDP3 package ^22^. No evidence for recombination was detected by three or more methods under a Bonferroni corrected *p*-value<0.001. Prior to estimation of final phylogenies, HA alignments were edited to remove the cleavage site and NA alignments were edited to remove hypervariable regions with poor alignment confidence. Trees were estimated from the sub-sampled alignments using RAxML under the GTR+G model and bootstrapped with 100 replicates ^21^. Accession numbers and alignment sizes are provided in Supplementary File 1. The varying sizes and/or missing HP clusters in some alignments are due to the lack of complete genomes for all strains in the public database.

### 3. Detecting HAPMs occurring across HP clusters

To screen for HAPMs within AIV genomes, all mutations (deletions, insertions and non-synonymous variable sites) present in ≥1% of the sequences in each sub-sampled alignment were coded as a discrete taxon trait and subjected to ancestral state reconstruction, using the maximum likelihood framework implemented in the “*ape*” R package ^23^. The resulting phylogenetic reconstructions were inspected visually to identify HAPMs shared by multiple HP clusters. HAPMs were identified if: (i) they occurred within 2 or more HP clusters, (ii) they were present in >60% of the sequences within the HP clusters, and (iii) they were not observed on internal branches anywhere else in the phylogeny (Supplementary File 2).

To illustrate in detail the evolutionary association between HAPMs and AIV HP clusters, we selected three HAPMS for each virus subtype for ancestral state reconstruction using Bayesian molecular clock phylogenetic analysis. These six examples were chosen to illustrate the range of evolutionary dynamics exhibited by HAPMs, and because they are all significantly associated with the HP phenotype (see next section). For each site, we estimated the posterior probability of amino acid states at tree nodes ancestral to HP clusters in the subsampled alignments, using the discrete trait evolution model in BEAST v1.8.4 ^24^. Time-scaled phylogenies were estimated under a SRD06 nucleotide substitution model, a strict molecular clock model and a Bayesian Skyline coalescent tree prior. Partitions were generated for each site, and ancestral states/transition rates were estimated for each mutation using an asymmetric continuous time Markov chain model ^25,26^. Two MCMC runs were computed for 100×10^6^ states or until convergence was reached. Maximum clade credibility (MCC) trees were summarized using TreeAnnotator, after removing 10% of the runs as burn-in ^24^. Additionally, we re-estimated the full evolutionary history of these HAPMs by mapping the amino acid changes onto the large-scale ML phylogenies generated in step 1 (Supplementary File 3).

### 4. Testing for genotype-to-phenotype associations

We developed a new evolutionary model to statistically test for associations between binary phenotypic traits and amino acid states in a phylogenetically-informed manner. We used this model to determine which of the HAPMs identified above were significantly associated with the HP phenotype. Our method serves the same purpose as a χ^2^ test, but correctly incorporates the phylogenetic history of the trait and phenotype of interest, as it is known that failing to account for these phylogenetic correlations can lead to erroneous false positives ^19,27^. The model assumes that phenotypes and amino acid states are discrete traits that change along phylogeny branches. Given an alignment, its phylogeny, and a set of binary values for each sequence, the method proceeds to test if a specific amino acid state is associated with the given phenotype. This is repeated for all amino acid states and all codons. A coupling parameter λ is used to measure the strength of association between an amino acid state and the phenotype state. The null hypothesis of zero association is represented by λ = 1, and can be rejected in favour of a positive association (λ >1) using a likelihood ratio test (LRT). The type I and II error rates of this test were evaluated using simulations. A full mathematical description of the model and simulation details are provided in Supplementary Text 5. We applied this model to our subsampled H5NX and H7NX alignments. The H5NX alignments were further reduced by collapsing clade C8 to only its most basal sequences, in order to avoid the overrepresentation of HP sequences.

### 5. Testing for diversifying positive selection

To identify positively-selected HAPMs, we used various *dN/dS* (ω) ratio estimation methods. The branch-site model A (BSA) in CODEML ^28^ was used to identify positively-selected sites (PSS) associated with specific phylogeny branches, namely the HP clusters. The statistical significance of the BSA model was compared to a null hypothesis with an ω=1 and evaluated using a LRT. The datamonkey server ^29^ was also used to test for sites evolving under pervasive and episodic positive selection under both site-(MEME: Mixed Effects Model of Evolution; SLAC: Single-Likelihood Ancestor Counting and FEL: Fixed Effects Likelihood) and branch-site models (BSREL: branch-site random effects likelihood) ^30,31,32,33^. Test significances were assessed using a LRT and subjected to Holm-Bonferroni multiple test correction. Finally, PSS were compared to the HAPMs detected and to the DEPS algorithm results (Directional Evolution of Protein Sequences using Maximum Likelihood) ^34^ (Supplementary Table 1). The LRT results and lists of the PSSs identified are provided in the Supplementary File 4.

### 6. Inferring mutation effects on protein stability and function

We undertook structural analyses to assess if the identified HAPMs might be functionally relevant. To determine the location and potential functional implications of the different HAPMs, we modelled protein 3D structures from existing structural data in the PDB database (https://pdb101.rcsb.org/), using representative HP sequences from our datasets as queries. For H7, we used the crystal structure of A/H7N2/New York/107/2003 AIV (PDB ID:3M5G), whilst for H5 we used the crystal structure of A/Vietnam/30262III/04 AIV (PDB ID: 2IBX). The high-resolution structure PDB 2Q06 was used to model NS1, and PDB 2Z16 was used to model M1. For the PB2/PB1/PA complex, the high-resolution structure from the bat Influenza A/H17N10 virus (PDB 4WSB) was used as a template, as this is the only structural data currently available for the complete influenza A virus polymerase complex ^35^. For the HA, M1, and NS1 proteins, we used the I-TASSER server ^36^ to generate models with the mutations of interest, that were compared manually to the PDB files for validation. For the polymerase complex, mutations were introduced to the superimposed 3D protein structures using PyMOL (Schrödinger, LLC). Finally, to predict which HAPMs had a possible effect on protein stability, we used the PIPS (Phylogenetic Inference of Protein Stability) program ^37^. This program implements an informed Bayesian approach and estimates, for each amino acid state, a value of free energy change (ddG) in terms of the thermodynamic stability of an unfolding/refolding protein, weighted under a phylogenetic background. Negative ddG values are predicted to be stabilizing for protein structure. Values close to zero are predicted to have little effect or to be only weakly stabilizing, whilst positive values are predicted to destabilize protein structure ^37^.

## RESULTS

### Large-scale phylogenetic analysis of independent HP clusters

Large-scale phylogenetic analysis confirmed that H7NX viruses are split into two major clades, the Eurasian and North American lineages (Supplementary Figure 1) ^38^. H7 HP outbreaks occurred independently through time and space and correspond to distinct lineages that, in most cases, display a unique pCS motif ^5,6^ (Figure 2). LP H7NX viruses carry a consensus amino acid sequence -PEXPKGRGLF-at the HA cleavage site, whereas a minimal consensus of -PEXPKXnK/RnRGLF-is identified for HP viruses (Figure 2a) ^39 11^. Nine H7NX well-supported clusters and spatio-temporally distinct HP lineages were identified in the HA phylogeny (Figure 2b) and in trees of other genome segments (Supplementary Figs. 2,3). Minor inconsistencies were observed for cluster C1, which segregated into C1.1 and C1.2 in the PB2, PB1, PA, M, and NS1 trees (Supplementary Figure 2).

**Figure 2.**
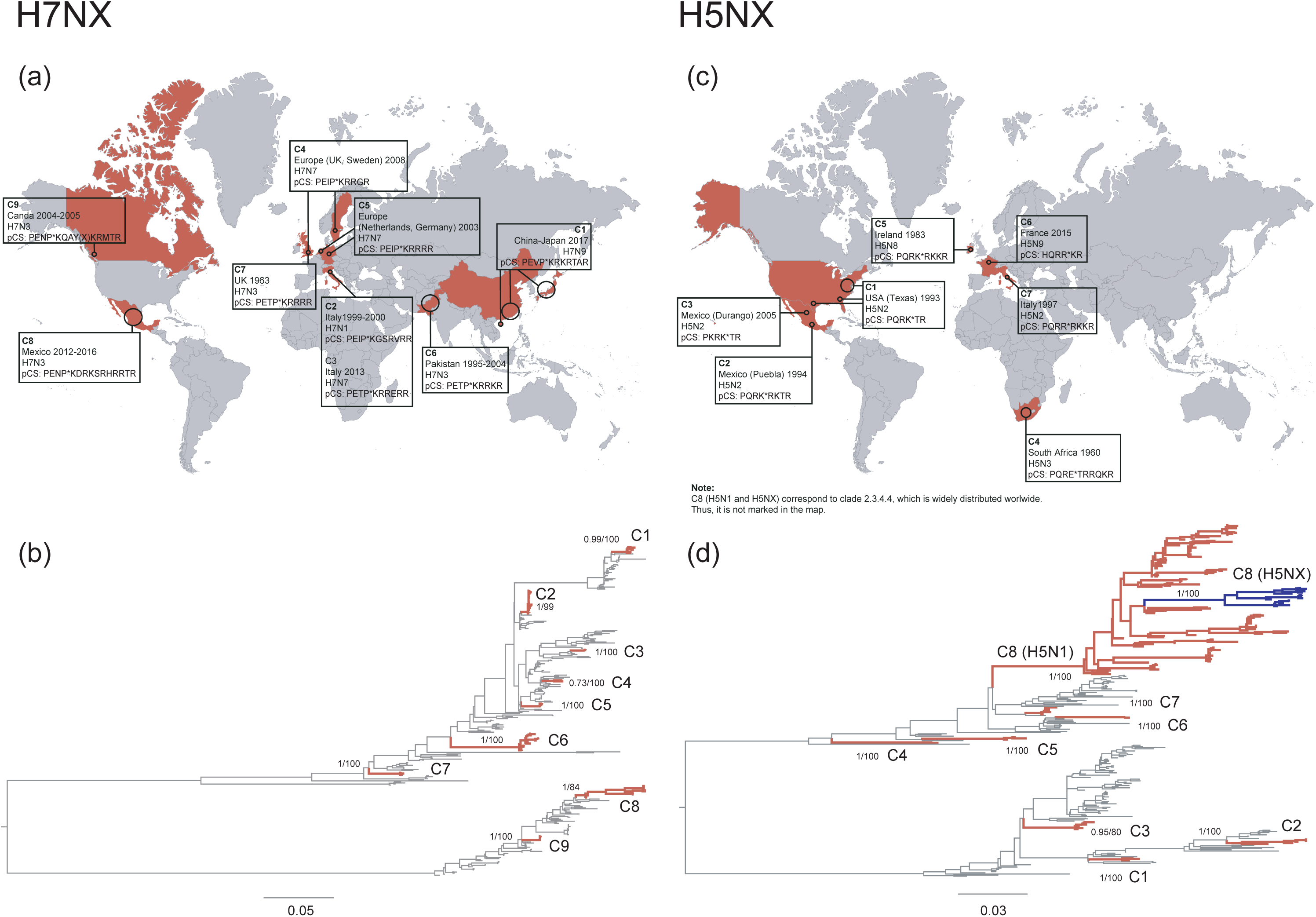
Independent spatio-temporal emergence of the HP phenotype. (a) Geographical occurrence of H7NX HP outbreaks. The year of circulation, virus subtype and consensus sequence for the polybasic cleavage site (pCS) are shown for the HP clusters analysed in this work. Countries of emergence are in red. (b) Phylogenetic distribution of HP outbreaks in the H7NX phylogeny. Distinct well-supported HP clusters within the HA tree are highlighted in red and named (C1-C9). Posterior probabilities and bootstrap support values are shown for nodes of interest. (c) Geographical occurrence of H5NX HP outbreaks, excluding C8 (clade 2.2) which is distributed worldwide. (d) Phylogenetic distribution of HP outbreaks in the H5NX phylogeny. Distinct well-supported HP clusters within the HA tree are highlighted in red and named (C1-C8). The H5NX HP viruses within cluster C8 (clade 2.2) are highlighted in blue.

The large-scale phylogeny of H5NX viruses shows a basal clade of HP and LP H5NX viruses circulating in wild birds worldwide from the late 1960s to the early 2000s (Figure 2d, Supplementary Figure 1). This basal clade is ancestral to the large and long-lived HP H5N1 lineage (named C8 here). This clade emerged in 2003 in China and subsequently spread to Asia, Africa, Europe and America ^1^ and persists until today ^5^. Within C8, there is a distinct group of HP H5NX viruses (H5N2, H5N6 and H5N8 subtypes) that circulates in both wild and domestic birds (Figure 2d). Eight H5NX HP clusters were identified as being spatio-temporally distinct and well-supported (Figure 2c). Most HP clusters displayed a unique pCS sequence, except for C8, reflecting the ongoing evolution of this long-lived lineage. As with the H7NX viruses, phylogenetic clustering of HP H5NX viruses was conserved within other genome segment phylogenies (Supplementary Figures 3,4).

### Identifying HAPMs significantly associated with the HP phenotype

The number of amino acid sites that were polymorphic (i.e. variants present at >1% frequency) varied among virus subtypes and genome segments. For H7NX viruses, the number of polymorphic sites per segment ranged from 21 (for M1) to 125 (for HA). Maximum likelihood ancestral state reconstructions revealed 16 parallel genetic changes (HAPMs) present in two or more H7NX HP lineages (Supplementary File 2). For H5NX viruses, between 12 (for M1) and 141 (for HA) polymorphic sites were detected for each segment. We identified 39 HAPMs present in two or more H5NX HP lineages (Supplementary File 2). Of these, 17 occurred within sub-lineages of cluster C8 and thus were not considered further. To focus attention on those HAPMs significantly associated with the HP phenotype, we developed a new phylogenetic model to test the null hypothesis of no association between mutations at each position and the HP phenotype (Supplementary Text 5). For H7NX viruses, we found a strong association with the HP phenotype for 9 out of 16 HAPMs (Table 1). For H5NX viruses, a strong association with the HP phenotype was supported for 12 out of 22 HAPMs (Table 1).

**TABLE 1.**
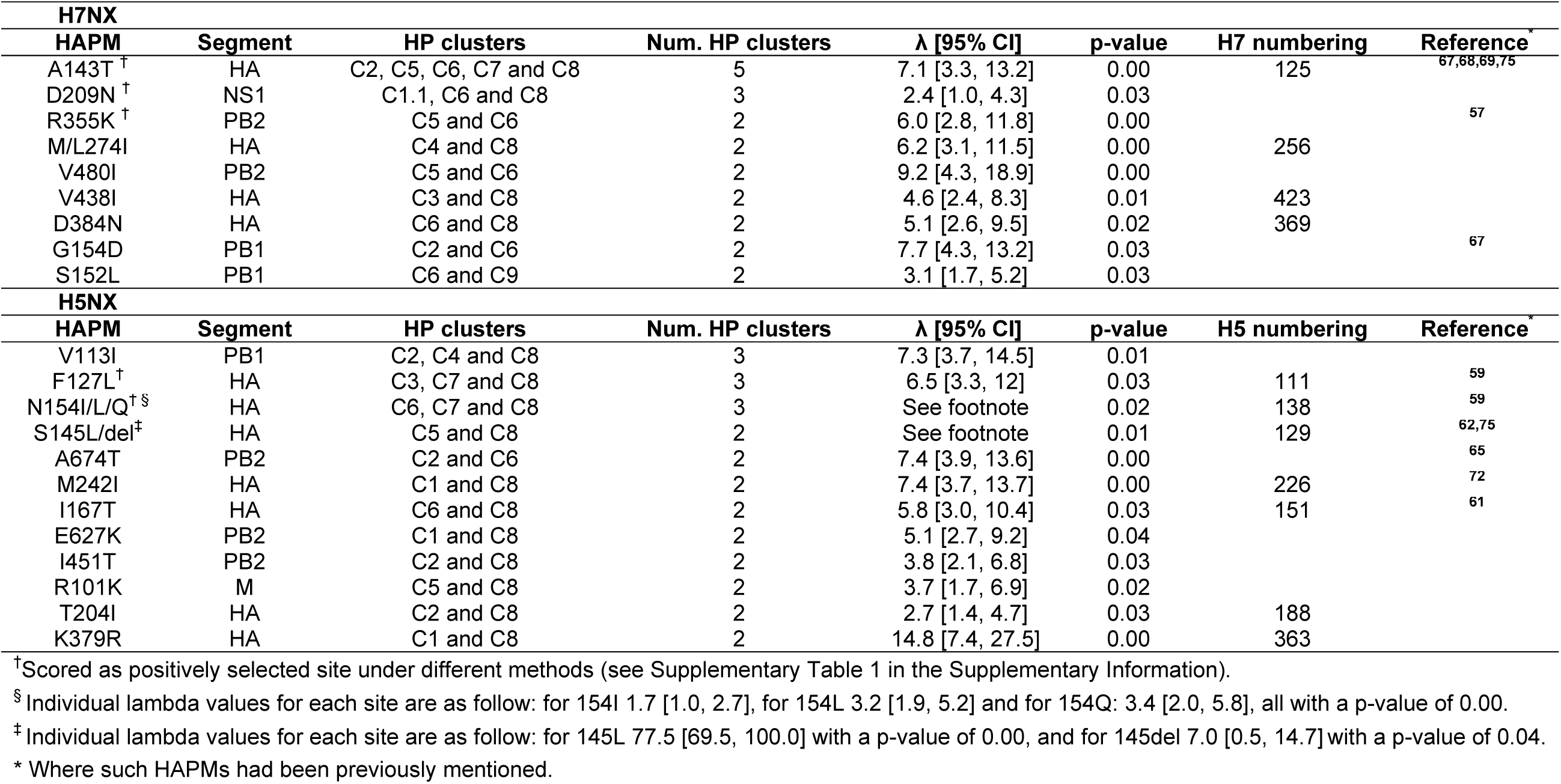
HAPMs significantly associated with the HP phenotype of H5 and H7 virus subtypes.

To illustrate the evolutionary dynamics of HAPMs, we chose a subset of six HAPMs that were significantly associated with the HP phenotype and subjected them to molecular clock phylogenetic analysis (Figure 3). We observed a variety of evolutionary patterns and in general obtained high posterior probability values for ancestral amino acids at nodes preceding the emergence of HP clusters. By way of illustration, consider site 355 of H7NX PB2. At this site K is present in HP clusters C5 and C6, but not in lineages representing the LP background (Figure 3a). A similar pattern is seen for mutation 627K in H5NX viruses (Figure 3f). More extensive parallel evolution is exhibited by mutations 143T in H7 (Figure 3b) and 127L in H5 (Figure 3d), which are shared by multiple HP clusters, but again not observed within LP lineages. Mutation 209N in H7NX NS1 is found within several HP clusters, but is also common in LP sequences of the American lineage (Figure 3c). Lastly, site 154 in H5 exhibits a more complex, step-wise evolutionary process, from N to I/L to Q, reflecting the transition from LP to HP (Figure 3e). N is present only in LP viruses, I/L occurs in HP viruses circulating prior to and during the early expansion of C8, and Q is only present in more recent HP viruses from C8 (Figure 3e).

**Figure 3.**
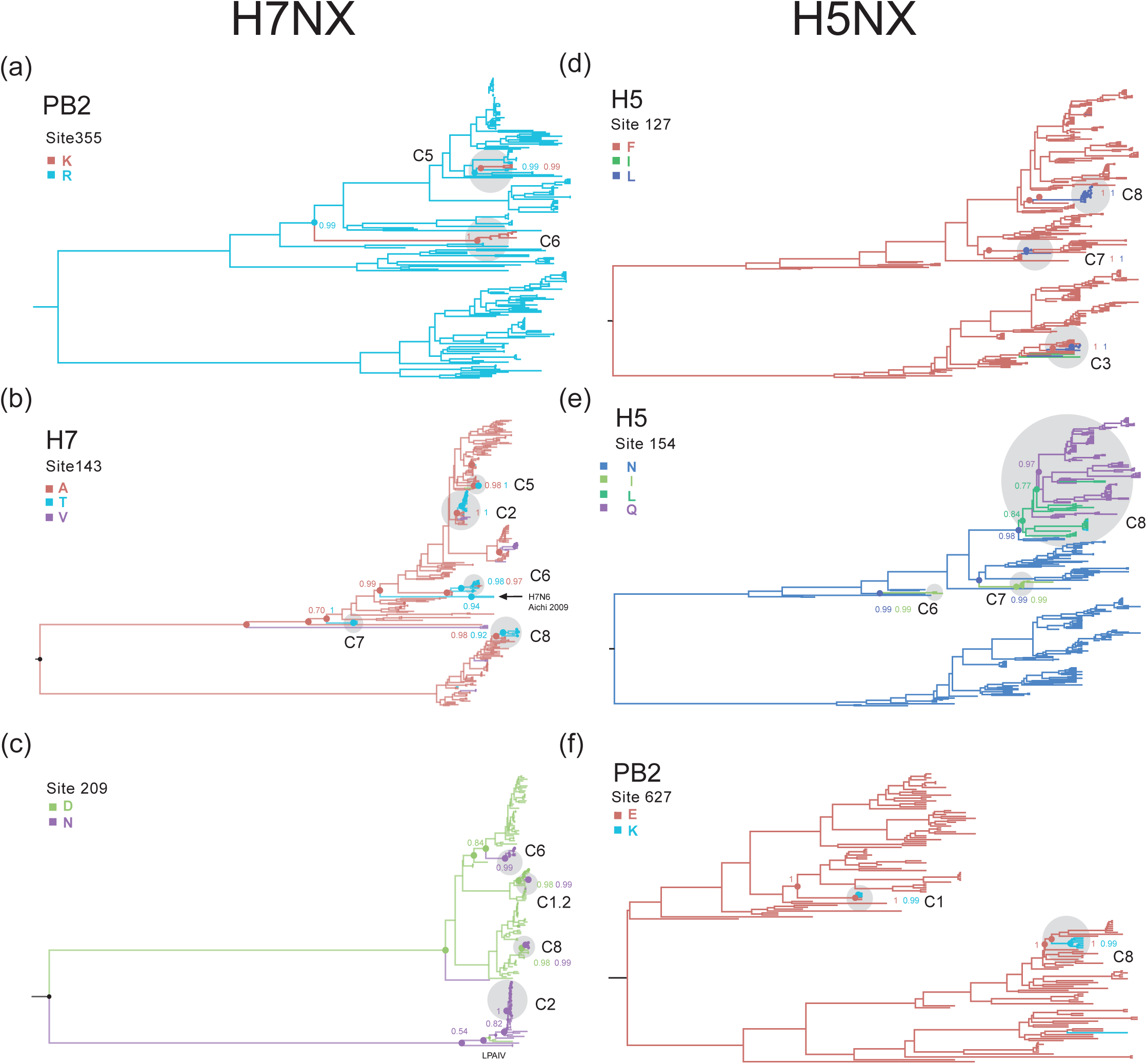
Time-scaled reconstruction of amino acid evolution at selected HAPMs. Maximum clade credibility trees are shown for six illustrative HAPMs (a-f) that exhibit a significant associated with the HP phenotype. Amino acid changes at each of the six sites were inferred using a Bayesian molecular clock phylogenetic approach. Branches are coloured according to the inferred amino acid states at internal nodes (tip states not shown). Ancestral nodes preceding the emergence of a given HAPM are highlighted by coloured circles. The posterior probabilities for the amino acid state occurring at specified node are shown. The HP clusters are highlighted by grey circles.

To facilitate computational analyses, we sub-sampled AIV alignments before detecting HAPMs (see Method step 2). Thus, the original alignments before subsampling may contain HP lineages that were not present after sub-sampling (Supplementary Figure 1). Thus, we investigated whether if these lineages also contained any HAPMs significantly associated with the HP phenotype, by mapping HAPM amino acid changes onto the full phylogenetic trees (Supplementary File 3). Since these HP lineages were not used to define HAPMs, mutations present in them constitute an independent test of our statistical approach. We found that, in addition to the HP clusters used here, mutation 154L in H5 was fixed in two independent HP H5N1 outbreaks in Vietnam from 2008-2013 ^40^, and in HP H5N2 viruses in China circulating from 2009-2015 ^41^ (Supplementary File 3). We also observed that mutation 143T is present in H7N6 viruses isolated from an outbreak in Aichi prefecture, Japan, in 2009 ^42^ (Figure 3b; Supplementary Files 1,3). Although this was not a HP outbreak, these viruses contained an additional arginine residue within the pCS (-PKGR to -PKRR), which triggered concerns about whether this lineage might evolve the HP phenotype. Fortunately, this lineage was successfully eradicated before any HP phenotype strains were observed ^42^.

Most HAPMs were not observed on phylogeny branches immediately ancestral to the HP clusters, suggesting that these appear shortly before, during, or after the emergence of the HP phenotype and making it difficult to discern if they are permissive or compensatory change. However mutation A143T in H7 is present in LP viral sequences that are basal and immediately ancestral to the emergence of the HP H7N1 outbreak in Italy 2000 ^43^. This observation supports the notion that at least some of the HAPMs identified here could be permissive.

### HAPMs associated with diversifying positive selection

Although parallel evolution is often caused by positive selection, parallel genetic changes can, in theory, also arise by chance ^14^. Consequently, we investigated whether if some HAPMs were inferred as being positively selected sites (PSS; see Methods section 5). Analyses under site- and branch-site models showed evidence for diversifying positive selection at some HAPMs, and for some HP clusters (H7NX clusters C4 and C7, and H5NX clusters C1, C2 and C4; Supplementary Table 1). As expected ^15^, *dN/dS* (ω) values for whole genome segments were <1 in all cases (data not shown), and the overall proportion of PSS detected was low (Supplementary Table 1, Supplementary File 4).

For the H7NX viruses, 4/16 HAMPs (corresponding to 4/9 of the HAPMs significantly associated with the HP phenotype) were identified as PSS. For the H5NX viruses, only 2/22 HAPMs (corresponding to 2/12 of the HAPMs significantly associated with the HP phenotype) were identified as PSS (see Supplementary Table 1 for details). At first impression, this suggests that most HAPMs significantly associated with the HP phenotype are not a direct consequence of positive selection, and may instead represent neutral variation that has arisen through genetic hitchhiking ^14,16^. However, if some HAPMs are selectively neutral in the absence of a pCS (LP phenotype), but offer a positive effect when a pCS is present (HP phenotype), then they may not be inferred as PSS using *dN/dS* methods yet still represent a permissive/compensatory change. This is a consequence of *dN/dS* estimation methods being conservative in detecting sites under positive selection in some circumstances ^44^. For example, mutation E627K in PB2, which was not inferred to be a PSS, is known to have a positive effect on the adaptation of AIV to mammalian hosts and increases replication efficacy ^45^. The role of selection on whole HP lineages is less clear: for example, experiments in turkey populations suggest that the HP and LP phenotypes of H7N1 viruses have indistinguishable transmission rates and basic reproductive numbers (R_0_) ^46^.

### Structural and protein stability analyses

Most of the HAPMs significantly associated with the HP phenotype had negative ddG values, as assessed under a Bayesian approach for inferring the stability effects of individual mutations from homologous protein sequences with a known phylogeny ^37^ (Figures 4 and 5). Thus, they are predicted to be fairly stabilizing/neutral changes to protein structure. One exception is E627K in H5NX PB2, which is predicted to be destabilizing. However, it has been shown that other compensatory mutations in PB2 appear to improve the genetic stability of E627K ^47^.

**Figure 4.**
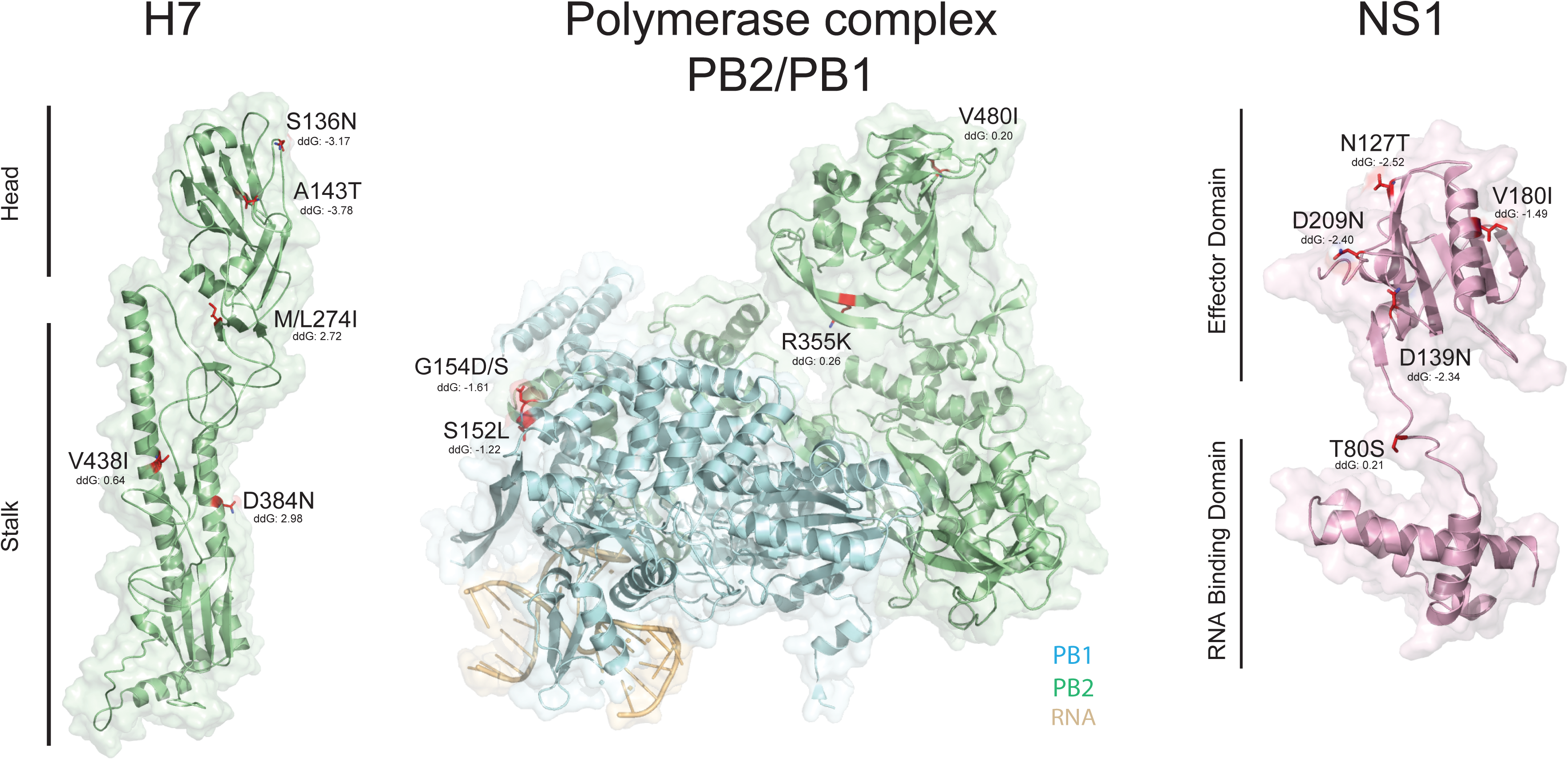
Functional relevance of HAPMs of H7NX viruses. Protein structure models for the HA (Hemagglutinin), PB2/PB1 (Polymerase complex) and the NS1 (Non-structural protein) of the H7NX viruses. Functionally relevant HAPMs are highlighted in red within the structural models; only those HAMPs that are significantly associated with the HP phenotype are indicated. Estimated values for free energy change (ddG) are indicated for these sites. Negative values or those close to zero can be interpreted as fairly stabilizing/neutral mutations with respect to protein structure.

**Figure 5.**
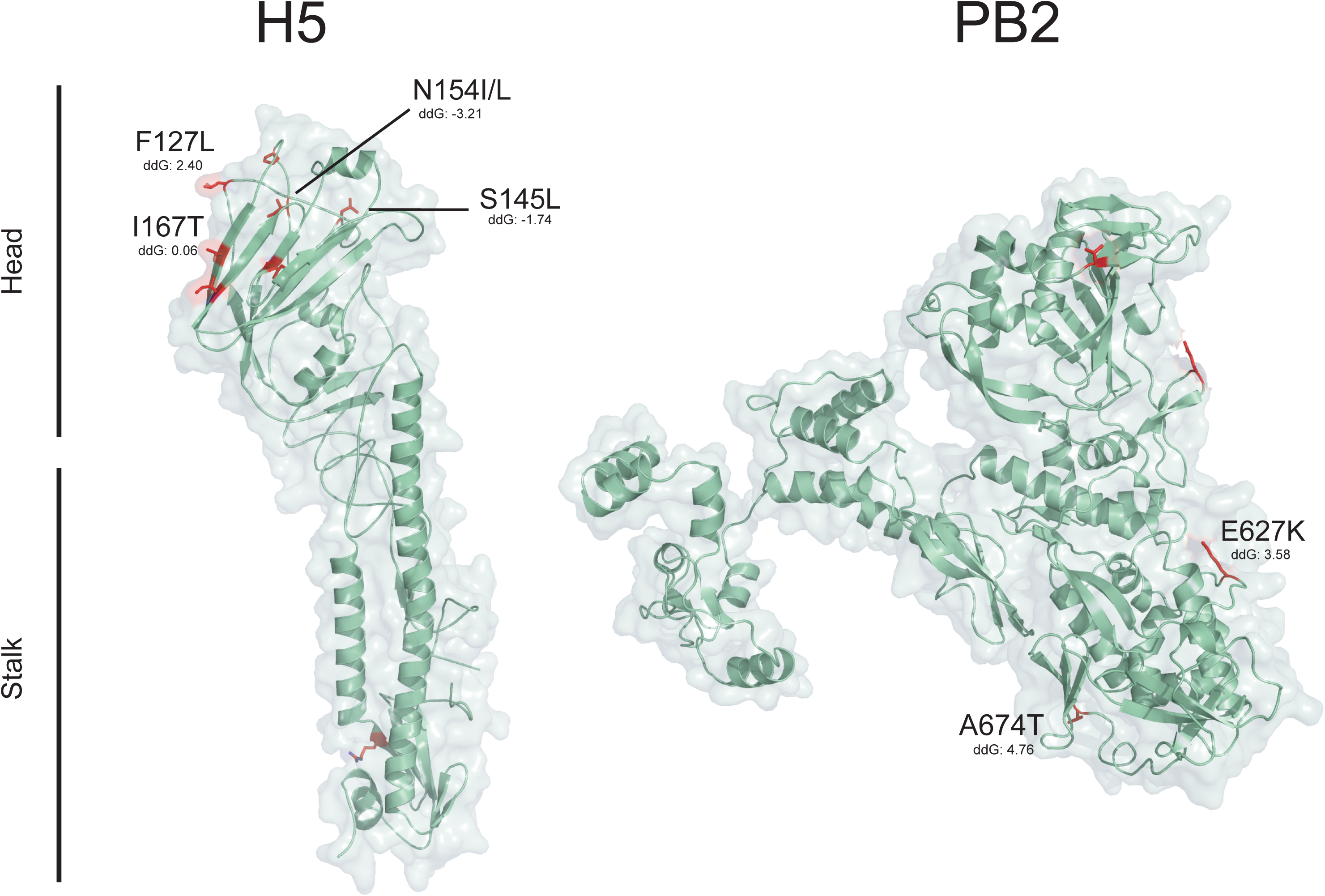
Functional relevance of HAPMs of H5NX viruses. Protein structure models for the HA (Hemagglutinin) and the PB2 polymerase subunit of the H5NX viruses. See Figure 4 legend for further details.

Structural analysis suggests that several H7NX HAPMs significantly associated with the HP are functionally relevant (Figure 4). A143T in HA is a non-conservative amino acid change located in a solvent-accessible region within the 130-loop proximal to the receptor surface. This is within antigenic pocket A, contributing to the electrostatic environment needed for ligand binding. Although it has no direct interaction with glycans ^48^, mutation A143T induces glycosylation of residue 141 by adding a N-linked glycosylation motif. This N-linked residue produces steric masking of the antigenic site and may affect HA folding and stability ^75^. Similarly, M/L271I in HA is located within the receptor binding subdomain ^49,50^. R355K in PB2 is a conservative amino acid change within a solvent-exposed loop of the electrostatic surface of the cap binding domain. This site is in direct contact with first phosphate after the cap ^51^ and correlates with high pathogenicity of AIVs in mouse models ^52^. D209N in NS1 is a conservative amino acid change located within a disordered tail interacting with importin alpha, a cellular host factor that promotes the entry of proteins to the cell nucleus (Figure 4), and may thus interfere with cell cycle regulation pathways ^53^.

HAMPs significantly associated with the HP phenotype in H5NX viruses were also found to be structurally relevant (Figure 5). Mutation F127L is a conservative amino acid change within the globular head of HA1, close to antigenic site 3 ^54^. N154I/L is a non-conservative amino acid change within antigenic site 2, and involves the loss of the asparagine-linked glycosylation site ^55^. Both these HAPMs correspond to variable sites related to escape mutants ^54^. However, they are not crucial for the protective efficacy of H5 vaccines, suggesting that the loss of the saccharide linkage may have little impact on antigenic variation ^50,55^. I167T is located in the surface-exposed region of the HA protein, and has been related to increased infectivity in the human lower respiratory tract in HP H5N1 viruses from Egypt ^56^. S145L in HA is also located in the surface exposed region and has been linked to increased virulence in poultry ^57^. Position 145 also undergoes occasional deletion (as observed in approximately 10% of sequences) and it has been argued that this deletion is critical for the antigenic evolution of long-lived HPAIV clusters ^75^. Mutation E627K in PB2 is a well-characterised adaptation marker associated with enhanced transmission in mammalian hosts ^58^, found frequently among H5NX viruses causing human infections ^7^. Site 627 is surface exposed and located on the “627 domain” and is essential for viral RNA replication and transcription in a cellular context ^59^ (Figure 5). Mutation A674T in PB2 is located at the interface of the polymerase complex trimer and is predicted to increase strength of the PB2 protein structure. In *in vivo* mouse models this mutation is a prerequisite for the acquisition of E627K, as it stabilizes polymerase activity when co-occurring with 627K ^60^, supporting the notion that this HAPM acts as a permissive mutation.

## DISCUSSION

The occurrence of parallel molecular evolution in AIV genomes can be used as a starting point for the identification of mutations associated with the evolution of the HP phenotype. We report combined evolutionary and structural evidence to support the hypothesis that some AIV parallel mutations are significantly associated with the HP phenotype and may potentially act as permissive/compensatory changes to facilitate the evolution of HP viruses. Previous studies have noted parallel genetic changes in independent AIV lineages ^61^, but these were not linked to the emergence of the HP phenotype. Similarly, specific instances of parallel genetic changes associated with the transition from a LP to a HP phenotype giving rise to single H7N1 outbreaks in Italy have also been described ^43,62,63^. We provide the first systematic and statistically-rigorous screening of AIV mutations associated with HP outbreaks. We demonstrate the effectiveness of phylogenetically-informed hypothesis testing coupled with comparative evolutionary and structural approaches to identify potentially important mutations exhibiting parallel evolution in H5 and H7 AIV genomes.

Permissive/compensatory mutations associated with the HP phenotype are expected to be favoured under a HP genetic background, and therefore occur repeatedly in HP lineages. Mutations that occur at nodes preceding and coinciding with the emergence of pCS-defined HP clusters (*e.g.* those that are present in the immediate LP ancestors and in the HP cluster), may represent permissive changes that facilitate the subsequent evolution of the HP phenotype. Conversely, mutations that occur only within HP clusters (after a pCS has arisen), may be more likely to be compensatory changes.

However, sampling of AIV genomes is likely biased towards HP outbreaks, resulting in a phylogenetic under-representation of the immediate LP ancestors of any given HP lineage. This under-sampling of local LP strains makes it difficult to discern whether the HAPMs correspond to permissive versus compensatory changes. Further molecular experiments are needed to understand how the parallel mutations detected here actually contribute to the evolution of the HP phenotype by, for example, stabilizing protein structure, altering host-virus or virus-virus inter-molecular interactions, or by combining additively to increase virulence. However, such experimental validation may take several years to complete especially if HP AIV gain-of-function experiments will be required ^64^. In the meantime, we should continue to exploit fully the information revealed by parallel molecular evolution, which represents a “natural” replicated experiment. This situation further highlights the importance of systematic sampling and whole genome sequencing of both HP viruses *and* their immediate LP ancestors, prior and during AIV outbreaks.

Some of the HAPMs identified here were previously identified either *in vitro* or *in silico* as being associated with HP AIV (Table 1) ^47,52,56-58,60^. Although this supports our results, it is possible that the number, biological roles, and genomic distribution of mutations involved in HP adaptation is currently underestimated. Notably, the HAPMs described here are not shared among the H7NX and H5NX virus subtypes, yet they commonly occur within the same genome segments (Table 1), suggesting that the evolution of the HP phenotype may interact with other well-studied biological processes, such as antigenic escape (determined by the HA protein), host-response modulation (regulated by PB2 and NS1) and replication efficiency (PB2). Our findings are in agreement with previous studies showing that the acquisition of a pCS may require coadaptation of the HA and other viral proteins ^65^.

Some of the mutations occurring close to antigenic sites may be indirectly associated with differential patterns of immunity in varied host populations, defined by complex factors such as animal age and sex distributions, previous exposure to infections/vaccines, or differences in host genetic diversity ^66^. Interestingly, mathematical modelling has suggested that the sporadic population dynamics of HP strains may be determined, at least in part, by their linkage to one or more specific antigenic variants, whose frequencies are driven by the dynamics of host immunity, rather than by mutability ^67,68^. Furthermore, the emergence of the HP phenotype has been often associated with virus adaptation from wild to domestic bird host populations ^61^. The strength of this association is difficult to assess in nature given the substantial bias (in both the literature and sequence databases) towards AIV infections and HP outbreaks in poultry versus those in wild birds. However, using our new model of mutation-trait association (Methods section 4; Supplementary Text 5), we found no association between any of the detected HAMPs (or any other mutation) and the host species (domestic versus wild bird; data not shown). This suggests that the parallel mutations identified here are not directly linked to transmission among domestic bird populations, although high population densities together with a limited genetic variability within the host (as occurs in intense farming facilities), may facilitate virus transmission. If acquiring a pCS does indeed reduce AIV transmission fitness in natural systems ^46^, then it is interesting to speculate that both intensive farming and compensatory/permissive mutations may act to facilitate the spread of a HP strain; the latter by increasing R_0_ and the former by lowering the threshold R_0_ required to sustain transmission in a population.

Lastly, we propose that the set of HAPMs associated with HPAIV lineages identified here (or any improvement to this set obtained in the future), may, with sufficient validation and testing, function as an early detection system capable of highlighting lineages that pose a greater risk of evolving the HP phenotype in the future. Importantly, the mutational panel identified here could potentially benefit to AIV control and bird health even if the mechanistic and functional actions of the mutations described are unknown. We suggest that circulating AIV strains in domestic and wild bird populations that do not contain a pCS site, but which exhibit a combination of the HAPMs identified here, are potential targets for more focussed and targeted genomic and epidemiological surveillance.

## AUTHOR CONTRIBUTIONS

MEZ and OGP designed research. MEZ performed research. MEZ and OGP analysed data. MG developed the statistical model and analysed data. BG and JT analysed genomic data. JRK and LC analysed structural data. TAB supervised structural data analysis. MEZ wrote the manuscript, with comments from all authors.

## ACKNOWLEDGEMENTS

MEZ is supported by an EMBO Long Term Fellowship (ALTF376-2017). MG, JT and OGP are supported by the EU’s Seventh Framework Programme (FP7/2007-2013)/European Research Council (614725-PATHPHYLODYN). BG is supported by the 2017 Universities of Academic Excellence Scholarship Program of the Secretariat for Higher Education, Science, Technology and Innovation of the Republic of Ecuador (ARSEQ-BEC-003163-2017) and by Universidad San Francisco de Quito. JRK and LC are supported by the Wellcome Investigator award (200835/Z/16/Z). The Wellcome Centre for Human Genetics is supported by grant 203141/Z/16/Z. TAB is supported by the Medical Research Council (MR/L009528/1).

## REFERENCES

1 Webster, R. G., Peiris, M., Chen, H. & Guan, Y. H5N1 outbreaks and enzootic influenza. Emerg Infect Dis 12, 3–8, doi:10.3201/eid1201.051024 (2006).

2 Buxton Bridges, C. et al. Risk of influenza A (H5N1) infection among health care workers exposed to patients with influenza A (H5N1), Hong Kong. J Infect Dis 181, 344–348, doi:10.1086/315213 (2000).

3 Du Ry van Beest Holle, M., Meijer, A., Koopmans, M. & de Jager, C. M. Human-to-human transmission of avian influenza A/H7N7, The Netherlands, 2003. Euro Surveill 10, 264–268 (2005).

4 Lam, T. T. & Pybus, O. G. Genomic surveillance of avian-origin influenza A viruses causing human disease. Genome Med 10, 50, doi:10.1186/s13073-018-0560-3 (2018).

5 Dhingra, M. S. et al. Geographical and Historical Patterns in the Emergences of Novel Highly Pathogenic Avian Influenza (HPAI) H5 and H7 Viruses in Poultry. Front Vet Sci 5, 84, doi:10.3389/fvets.2018.00084 (2018).

6 Short, K. R. et al. One health, multiple challenges: The inter-species transmission of influenza A virus. One Health 1, 1–13, doi:10.1016/j.onehlt.2015.03.001 (2015).

7 Hatta, M., Gao, P., Halfmann, P. & Kawaoka, Y. Molecular basis for high virulence of Hong Kong H5N1 influenza A viruses. Science 293, 1840–1842, doi:10.1126/science.1062882 (2001).

8 Jackson, D., Hossain, M. J., Hickman, D., Perez, D. R. & Lamb, R. A. A new influenza virus virulence determinant: the NS1 protein four C-terminal residues modulate pathogenicity. Proc Natl Acad Sci U S A 105, 4381–4386, doi:10.1073/pnas.0800482105 (2008).

9 Schmolke, M. et al. Differential contribution of PB1-F2 to the virulence of highly pathogenic H5N1 influenza A virus in mammalian and avian species. PLoS Pathog 7, e1002186, doi:10.1371/journal.ppat.1002186 (2011).

10 Horimoto, T. & Kawaoka, Y. Reverse genetics provides direct evidence for a correlation of hemagglutinin cleavability and virulence of an avian influenza A virus. J Virol 68, 3120–3128 (1994).

11 Abdelwhab, E. M. et al. Composition of the Hemagglutinin Polybasic Proteolytic Cleavage Motif Mediates Variable Virulence of H7N7 Avian Influenza Viruses. Sci Rep 6, 39505, doi:10.1038/srep39505 (2016).

12 OFFLU. OIE/FAO Network of expertise on animal influenza: Influenza A Cleavage Sites Report 2018, <http://www.offlu.net/fileadmin/home/en/resource-centre/pdf/Influenza_A_Cleavage_Sites.pdf > (2018).

13 Stern, D. L. The genetic causes of convergent evolution. Nat Rev Genet 14, 751–764, doi:10.1038/nrg3483 (2013).

14 Gutierrez, B., Escalera-Zamudio, M. & Pybus, O. G. Parallel molecular evolution and adaptation in viruses. Current Opinion in Virology 34, 90–96, doi:10.1016/j.coviro.2018.12.006. (2019).

15 Pond, S. L., Murrell, B. & Poon, A. F. Evolution of viral genomes: interplay between selection, recombination, and other forces. Methods Mol Biol 856, 239–272, doi:10.1007/978-1-61779-585-5_10 (2012).

16 Frost, S. D. W., Magalis, B. R. & Kosakovsky Pond, S. L. Neutral Theory and Rapidly Evolving Viral Pathogens. Mol Biol Evol 35, 1348–1354, doi:10.1093/molbev/msy088 (2018).

17 Bloom, J. D., Gong, L. I. & Baltimore, D. Permissive secondary mutations enable the evolution of influenza oseltamivir resistance. Science 328, 1272–1275, doi:10.1126/science.1187816 (2010).

18 Nao, N. et al. Genetic Predisposition To Acquire a Polybasic Cleavage Site for Highly Pathogenic Avian Influenza Virus Hemagglutinin. MBio 8, doi:10.1128/mBio.02298-16 (2017).

19 Bhattacharya, T. et al. Founder effects in the assessment of HIV polymorphisms and HLA allele associations. Science 315, 1583–1586, doi:10.1126/science.1131528 (2007).

20 Katoh, K., Misawa, K., Kuma, K. & Miyata, T. MAFFT: a novel method for rapid multiple sequence alignment based on fast Fourier transform. Nucleic Acids Res 30, 3059–3066 (2002).

21 Stamatakis, A. RAxML version 8: a tool for phylogenetic analysis and post-analysis of large phylogenies. Bioinformatics 30, 1312–1313, doi:10.1093/bioinformatics/btu033 (2014).

22 Martin, D. P. et al. RDP3: a flexible and fast computer program for analyzing recombination. Bioinformatics 26, 2462–2463, doi:10.1093/bioinformatics/btq467 (2010).

23 Paradis, E. & Schliep, K. ape 5.0: an environment for modern phylogenetics and evolutionary analyses in R. Bioinformatics, doi:10.1093/bioinformatics/bty633 (2018).

24 Drummond, A. J., Suchard, M. A., Xie, D. & Rambaut, A. Bayesian phylogenetics with BEAUti and the BEAST 1.7. Mol Biol Evol 29, 1969–1973, doi:10.1093/molbev/mss075 (2012).

25 Ferreira, M. A. R. & Suchard, M. A. Bayesian analysis of elapsed times in continuous-time Markov chains. Canadian Journal of Statistics 36, 355–368, doi:https://dx.doi.org/10.1002/cjs.5550360302 (2008).

26 Lemey, P., Rambaut, A., Drummond, A. J. & Suchard, M. A. Bayesian phylogeography finds its roots. PLoS Comput Biol 5, e1000520, doi:10.1371/journal.pcbi.1000520 (2009).

27 Harvey, P. H. & Pagel, M. D. The Comparative Method in Evolutionary Biology. (Oxford University Press, 1991).

28 Yang, Z. PAML 4: phylogenetic analysis by maximum likelihood. Mol Biol Evol 24, 1586–1591, doi:10.1093/molbev/msm088 (2007).

29 Delport, W., Poon, A. F., Frost, S. D. & Kosakovsky Pond, S. L. Datamonkey 2010: a suite of phylogenetic analysis tools for evolutionary biology. Bioinformatics 26, 2455–2457, doi:10.1093/bioinformatics/btq429 (2010).

30 Kosakovsky Pond, S. L. & Frost, S. D. Not so different after all: a comparison of methods for detecting amino acid sites under selection. Mol Biol Evol 22, 1208–1222, doi:10.1093/molbev/msi105 (2005).

31 Murrell, B. et al. Detecting individual sites subject to episodic diversifying selection. PLoS Genet 8, e1002764, doi:10.1371/journal.pgen.1002764 (2012).

32 Murrell, B. et al. Gene-wide identification of episodic selection. Mol Biol Evol 32, 1365–1371, doi:10.1093/molbev/msv035 (2015).

33 Smith, M. D. et al. Less is more: an adaptive branch-site random effects model for efficient detection of episodic diversifying selection. Mol Biol Evol 32, 1342–1353, doi:10.1093/molbev/msv022 (2015).

34 Kosakovsky Pond, S. L., Poon, A. F., Leigh Brown, A. J. & Frost, S. D. A maximum likelihood method for detecting directional evolution in protein sequences and its application to influenza A virus. Mol Biol Evol 25, 1809–1824, doi:10.1093/molbev/msn123 (2008).

35 Reich, S. et al. Structural insight into cap-snatching and RNA synthesis by influenza polymerase. Nature 516, 361-+, doi:10.1038/nature14009 (2014).

36 Yang, J. et al. The I-TASSER Suite: protein structure and function prediction. Nat Methods 12, 7–8, doi:10.1038/nmeth.3213 (2015).

37 Bloom, J. D. & Glassman, M. J. Inferring stabilizing mutations from protein phylogenies: application to influenza hemagglutinin. PLoS Comput Biol 5, e1000349, doi:10.1371/journal.pcbi.1000349 (2009).

38 Xu, Y. et al. Limited Antigenic Diversity in Contemporary H7 Avian-Origin Influenza A Viruses from North America. Sci Rep 6, 20688, doi:10.1038/srep20688 (2016).

39 Bosch, F. X., Garten, W., Klenk, H. D. & Rott, R. Proteolytic cleavage of influenza virus hemagglutinins: primary structure of the connecting peptide between HA1 and HA2 determines proteolytic cleavability and pathogenicity of Avian influenza viruses. Virology 113, 725–735 (1981).

40 Nguyen, T. et al. Characterization of a highly pathogenic avian influenza H5N1 virus sublineage in poultry seized at ports of entry into Vietnam. Virology 387, 250–256, doi:10.1016/j.virol.2009.03.006 (2009).

41 Zhao, G. et al. Novel reassortant highly pathogenic H5N2 avian influenza viruses in poultry in China. PLoS One 7, e46183, doi:10.1371/journal.pone.0046183 (2012).

42 Uchida, Y. et al. Genetic characterization and susceptibility on poultry and mammal of H7N6 subtype avian influenza virus isolated in Japan in 2009. Vet Microbiol 147, 1–10, doi:10.1016/j.vetmic.2010.05.037 (2011).

43 Monne, I. et al. Emergence of a highly pathogenic avian influenza virus from a low-pathogenic progenitor. J Virol 88, 4375–4388, doi:10.1128/JVI.03181-13 (2014).

44 Anisimova, M. & Kosiol, C. Investigating protein-coding sequence evolution with probabilistic codon substitution models. Mol Biol Evol 26, 255–271, doi:10.1093/molbev/msn232 (2009).

45 Linster, M. et al. Identification, characterization, and natural selection of mutations driving airborne transmission of A/H5N1 virus. Cell 157, 329–339, doi:10.1016/j.cell.2014.02.040 (2014).

46 Saenz, R. A. et al. Quantifying transmission of highly pathogenic and low pathogenicity H7N1 avian influenza in turkeys. PLoS One 7, e45059, doi:10.1371/journal.pone.0045059 (2012).

47 Long, J. S. et al. The effect of the PB2 mutation 627K on highly pathogenic H5N1 avian influenza virus is dependent on the virus lineage. J Virol 87, 9983–9996, doi:10.1128/JVI.01399-13 (2013).

48 Yang, H., Chen, L. M., Carney, P. J., Donis, R. O. & Stevens, J. Structures of receptor complexes of a North American H7N2 influenza hemagglutinin with a loop deletion in the receptor binding site. PLoS Pathog 6, e1001081, doi:10.1371/journal.ppat.1001081 (2010).

49 Lu., J. et al. Molecular evolution, diversity and adaptation of H7N9 influenza A viruses in China. Emerging Infectious Diseases 24, doi:DOI: 10.3201/eid2410.171063 (2018).

50 Ha, Y., Stevens, D. J., Skehel, J. J. & Wiley, D. C. H5 avian and H9 swine influenza virus haemagglutinin structures: possible origin of influenza subtypes. EMBO J 21, 865–875, doi:10.1093/emboj/21.5.865 (2002).

51 Guilligay, D. et al. The structural basis for cap binding by influenza virus polymerase subunit PB2. Nat Struct Mol Biol 15, 500–506, doi:10.1038/nsmb.1421 (2008).

52 Katz, J. M. et al. Molecular correlates of influenza A H5N1 virus pathogenesis in mice. J Virol 74, 10807–10810 (2000).

53 Melen, K. et al. Nuclear and nucleolar targeting of influenza A virus NS1 protein: striking differences between different virus subtypes. J Virol 81, 5995–6006, doi:10.1128/JVI.01714-06 (2007).

54 Stevens, J. et al. Structure and receptor specificity of the hemagglutinin from an H5N1 influenza virus. Science 312, 404–410, doi:10.1126/science.1124513 (2006).

55 Hoffmann, E., Lipatov, A. S., Webby, R. J., Govorkova, E. A. & Webster, R. G. Role of specific hemagglutinin amino acids in the immunogenicity and protection of H5N1 influenza virus vaccines. Proc Natl Acad Sci U S A 102, 12915–12920, doi:10.1073/pnas.0506416102 (2005).

56 Saleh, M. M., Selim, A., Shahera, A. A., El-Shahidy, M. S. & El-Trabilli, M. M. Molecular characterization of recently isolated highly pathogenic avian influenza virus (H5N1) in Egypt. Nature and Science 15 (2017).

57 Perdue, M. L. & Suarez, D. L. Structural features of the avian influenza virus hemagglutinin that influence virulence. Vet Microbiol 74, 77–86 (2000).

58 Steel, J., Lowen, A. C., Mubareka, S. & Palese, P. Transmission of influenza virus in a mammalian host is increased by PB2 amino acids 627K or 627E/701N. PLoS Pathog 5, e1000252, doi:10.1371/journal.ppat.1000252 (2009).

59 Nilsson, B. E., Te Velthuis, A. J. & Fodor, E. Role of the PB2 627 Domain in Influenza A Virus Polymerase Function. J Virol 91, doi:10.1128/JVI.02467-16 (2017).

60 Lee, C. Y. et al. Prerequisites for the acquisition of mammalian pathogenicity by influenza A virus with a prototypic avian PB2 gene. Sci Rep 7, 10205, doi:10.1038/s41598-017-09560-z (2017).

61 Lebarbenchon, C. & Stallknecht, D. E. Host shifts and molecular evolution of H7 avian influenza virus hemagglutinin. Virol J 8, 328, doi:10.1186/1743-422X-8-328 (2011).

62 Fusaro, A. et al. Evolutionary trajectories of two distinct avian influenza epidemics: Parallelisms and divergences. Infect Genet Evol 34, 457–466, doi:10.1016/j.meegid.2015.05.020 (2015).

63 Banks, J. et al. Changes in the haemagglutinin and the neuraminidase genes prior to the emergence of highly pathogenic H7N1 avian influenza viruses in Italy. Arch Virol 146, 963–973 (2001).

64 Casadevall, A. & Imperiale, M. J. Risks and benefits of gain-of-function experiments with pathogens of pandemic potential, such as influenza virus: a call for a science-based discussion. MBio 5, e01730–01714, doi:10.1128/mBio.01730-14 (2014).

65 Veits, J. et al. Avian influenza virus hemagglutinins H2, H4, H8, and H14 support a highly pathogenic phenotype. Proc Natl Acad Sci U S A 109, 2579–2584, doi:10.1073/pnas.1109397109 (2012).

66 Hill, S. C. et al. Antibody responses to avian influenza viruses in wild birds broaden with age. Proc Biol Sci 283, doi:10.1098/rspb.2016.2159 (2016).

67 Wikramaratna, P. S., Pybus, O. G. & Gupta, S. Contact between bird species of different lifespans can promote the emergence of highly pathogenic avian influenza strains. Proc Natl Acad Sci U S A 111, 10767–10772, doi:10.1073/pnas.1401849111 (2014).

68 Thompson, C. P. et al. A naturally protective epitope of limited variability as an influenza vaccine target. Nat Commun 9, 3859, doi:10.1038/s41467-018-06228-8 (2018).

