## Supplementary Information for "Parallel Evolution in the Emergence of Highly Pathogenic Avian Influenza A Viruses"

**Supplementary Table 1.** HAPMs and HP clusters inferred to be evolving under positive selection

**Supplementary Figure 1.** Large-sale phylogenetic trees for the HA genome segments of the H7NX and H5NX viruses

**Supplementary Figure 2.** Trees for the genome segments of the H7NX viruses

**Supplementary Figure 3.** Trees for the NA genome segments (N1-N9 subtypes) of H7NX and H5NX viruses

**Supplementary Figure 4.** Trees for the genome segments of the H5NX viruses

**Supplementary Text 5.** Full mathematical description of the model and simulations

**ADDITIONAL SUPPLEMENTARY FILES (available in DRYAD)**

**Supplementary File 1.** Accession numbers for reduced datasets

H7_ACCESSIONS.xls

H5_ACCESSIONS.xls

**Supplementary File 2.** ML trees for the reconstructions of ancestral states

RASML_H7.pdf

RASML_H5.pdf

**Supplementary File 3.** HA large-scale phylogenetic trees coloured according to amino acid states/ phenotype.

LARGE_SCALE_H7.pdf

LARGE_SCALE_H5.pdf

**Supplementary File 4.** Likelihood Ratio Tests and PSS scored under BST

H7_LRT_CODEML.xls

H7_PSS_CODEML_BST.xls

H5_LRT_CODEML.xls

H5_PSS_CODEML_BST.xls

#### Supplementary Table 1. HAPMs and HP clusters inferred to be evolving under positive selection

| **H7NX** | Site models (pervasive selection) | | | | Site models (episodic selection) | | Branch-site models (episodic selection) | | | | | | Directional Selection (DEPS) |
|---|---|---|---|---|---|---|---|---|---|---|---|---|---|
| Gene* | SLAC ≤ 0.05 | SITE * | FEL ≤ 0.05 | SITE * | MEME ≤ 0.05 | SITE * | BSA ≤ 0.05 [‡] | CLUSTER | SITE * [§] | bsREL ≤ 0.05 | CLUSTER | SITE | DEPS: Mutation/EBF value[‖] |
| PB2 | NO | -- | NO | -- | YES | -- | NO | C6, C7 | **355** | NO | -- | -- | -- |
| PB1 | NO | -- | YES | **152** | YES | -- | NO | C9 | **152** | NO | -- | -- | -- |
| PA | NO | -- | NO | -- | NO | -- | NO | -- | -- | NO | -- | -- | -- |
| H7 | NO | -- | YES | **143** | YES | -- | YES | **C4** | -- | YES | **C4, C7** | -- | **143T/1139** |
| NP | NO | -- | NO | -- | YES | -- | NO | -- | -- | NO | -- | -- | -- |
| N3 | NO | -- | YES | -- | NO | -- | NO | -- | -- | NO | -- | -- | -- |
| M | NO | -- | NO | -- | YES | -- | NO | -- | -- | NO | -- | -- | -- |
| NS | NO | -- | YES | **180, 209** | YES | **180, 209** | NO | C8 | **180** | NO | -- | -- | **209N/10E5** |
| **H5NX** | SLAC ≤ 0.05 | SITE * | FEL ≤ 0.05 | SITE * | MEME ≤ 0.05 | SITE * | BSA ≤ 0.05 [‡] | CLUSTER | SITE * [§] | bsREL ≤ 0.05 | CLUSTER | SITE | DEPS: Mutation/EBF value[‖] |
| PB2 | NO | -- | NO | -- | YES | -- | NO | -- | -- | YES | **C1 TEX** | -- | -- |
| PB1 | NO | -- | NO | -- | NO | -- | NO | -- | -- | NO | -- | -- | -- |
| PA | NO | -- | NO | -- | NO | -- | NO | -- | -- | YES | **C4 SA** | -- | -- |
| H5 | YES | -- | YES | **154** | YES | **127, 154** | NO | -- | -- | NO | -- | -- | **S145L/9814**, **N154I/101**, **S157P/>105**, **I167T/>105** |
| NP | NO | -- | NO | -- | NO | -- | NO | -- | -- | NO | -- | -- | -- |
| N2 | NO | -- | YES | -- | YES | -- | NO | -- | -- | NO | -- | -- | -- |
| M | NO | -- | NO | -- | YES | -- | NO | -- | -- | NO | -- | -- | -- |
| NS | YES | -- | YES | **213** | YES | **213** | YES | **C2 PUE** | -- | NO | -- | -- | **P213S/>105** |

[*] Genes/sites shown in this table correspond only to those where HAPMs were detected by reconstruction of ancestral states
[‡] Although the significance for the BSA test can be negative due to very small global proportion of PSS, sites can still be significantly scored under BEB
[§] PSS scored under BEB (PP ≥ 90%)
[‖] Empirical Bayes Factor (EBF) for convergent evolution using Directional Evolution of Protein Sequences (DEPS)

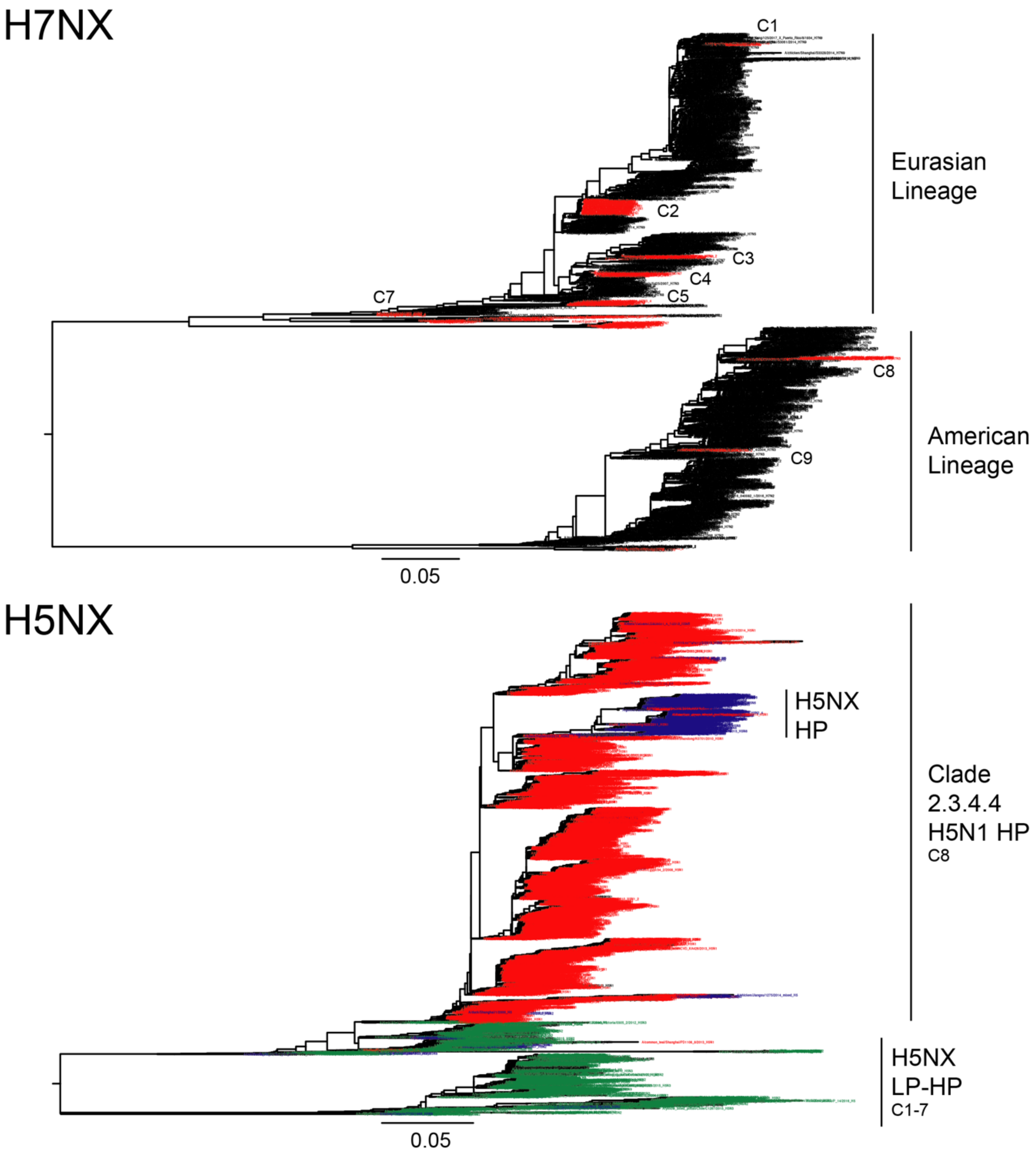


**Supplementary Figure 1. Large-sale phylogenetic trees for HA genome segments of H7NX and H5NX viruses**

For the H7NX tree (n = 2217), sequence names corresponding to highly pathogenic genotypes (HP) are colour-coded in red, whilst sequences corresponding to low-pathogenic genotypes are shown in black. For the H5NX tree (n = 5650), sequences corresponding to H5N1 HP genotypes are shown in red, whilst sequences in blue correspond to HP H5NX genotypes (excluding H5N1). Sequence names in green represent LP H5NX (excluding H5N1) viruses. For each virus subtype, the HP clusters used in this work are indicated.

# H7NX

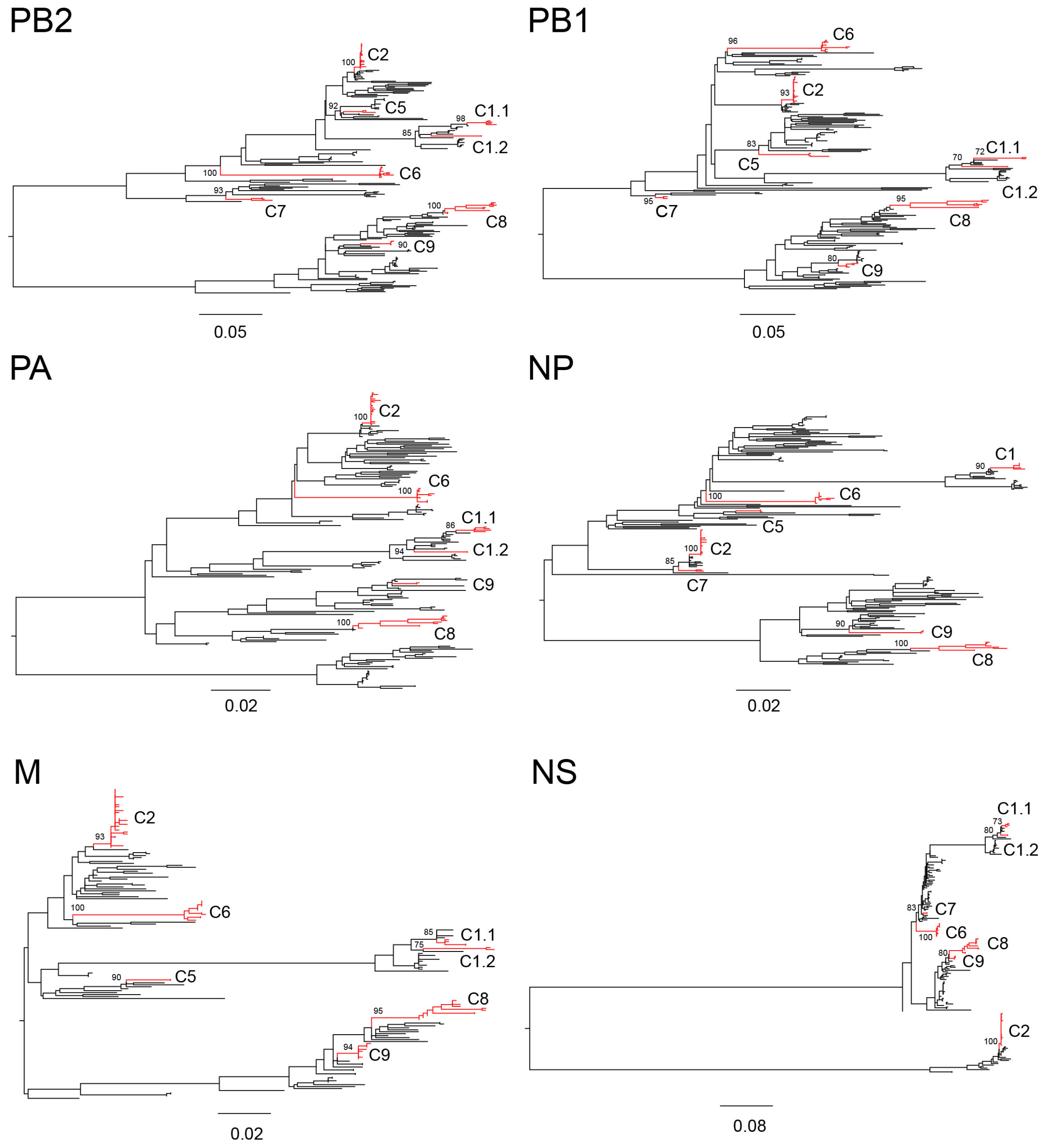


**Supplementary Figure 2. Maximum likelihood trees for the internal genome segments of H7NX viruses**

For H7NX viruses, the total numbers of sequences retrieved were: PB2=1856, PB1=1787, HA=2217, PA=1760, NP=1513, NA=1739, M1=1682 and NS1=1231. ML trees for the PB2, PB1, PA, NP, M1 and NS1 viral genome segments estimated using the sub-sampled alignments (described in Methods section 2, main text). No evidence of reassortment was observed for the HP lineages. Selected HP clusters (C1-C9) are highlighted in red and bootstrap values are shown for branches of interest. Minor inconsistencies among segments observed for cluster C1 include the segregation into C1.1 and C1.2 in the PB2, PB1, PA, M and NS1 trees. Clusters are missing in some of the phylogenies is due to the lack of corresponding sequences for all viral genome segments.

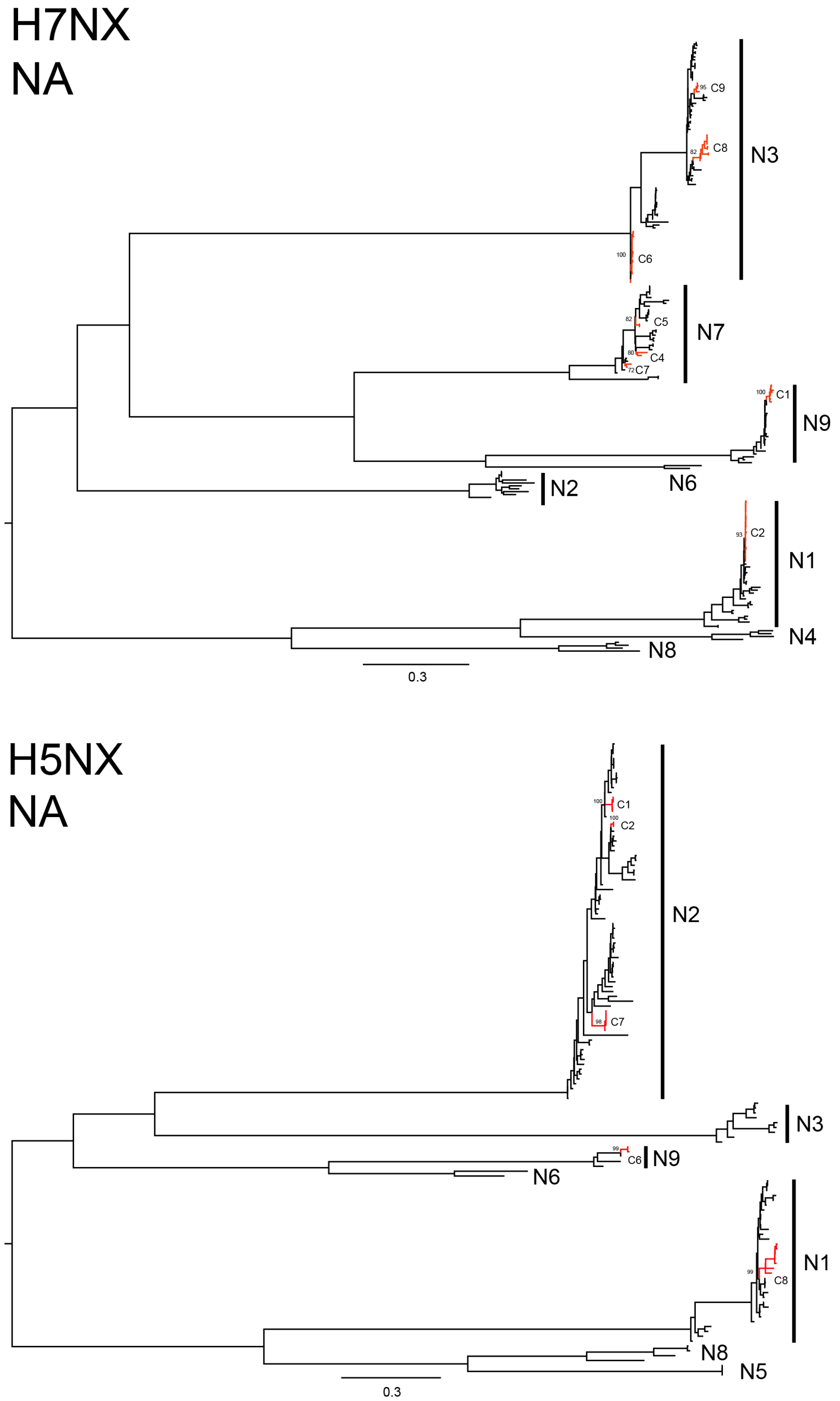


**Supplementary Figure 3. Maximum likelihood trees for the NA genome segments (N1-N9 subtypes) of H7NX and H5NX viruses**

ML trees for the NA genome segments of H7NX and H5NX viruses. HP clusters that occur in trees of different NA subtypes are indicated in red. Bootstrap values are shown for branches of interest.

# H5NX

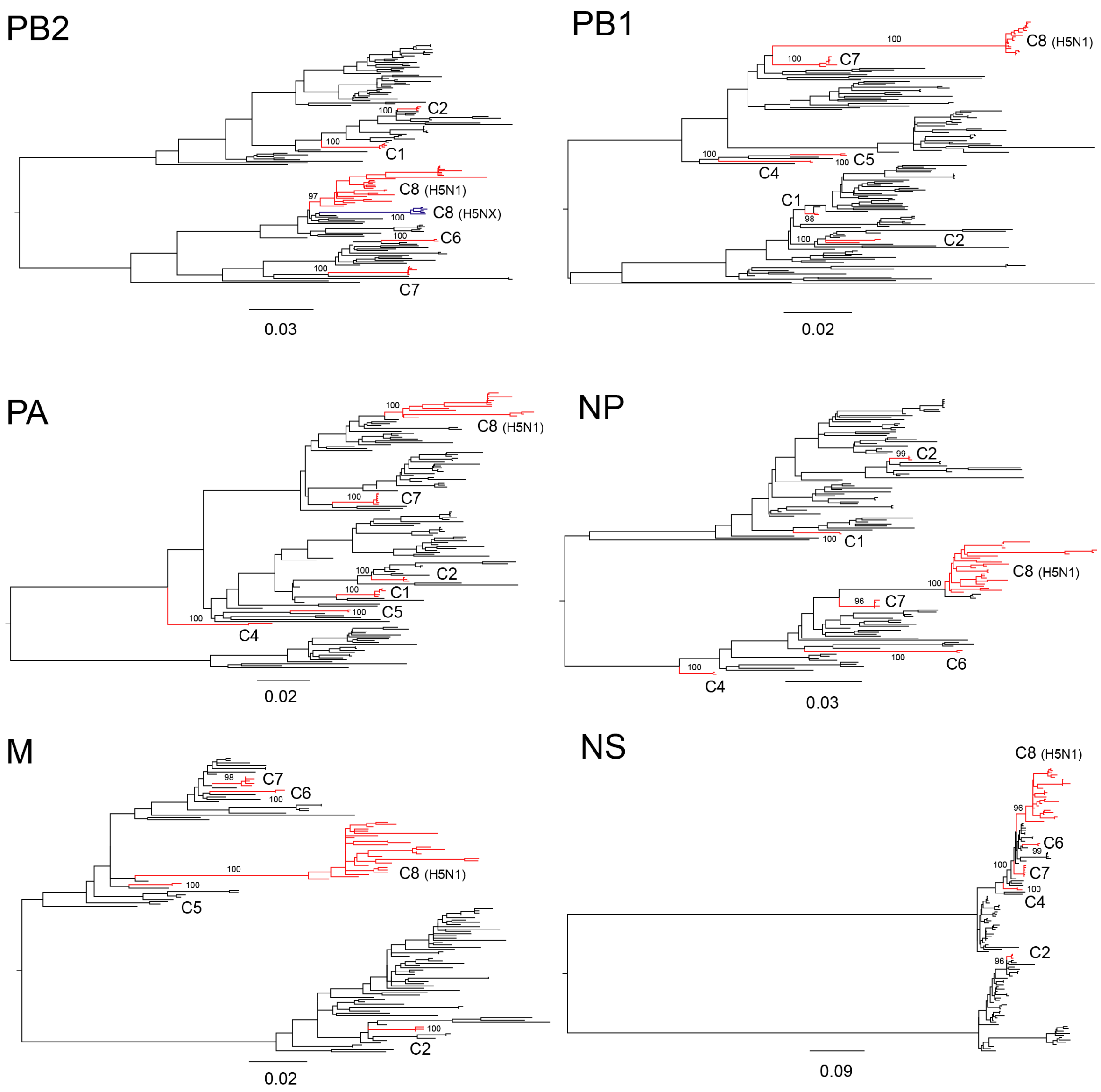


**Supplementary Figure 4. Maximum likelihood trees for the internal genome segments of H5NX viruses**

For H5NX viruses the numbers of sequences were: PB2=3554, PB1=3817, PA=3421, HA=5650, NP=3230, NA=4202, M1=2824 and NS1=3690. ML trees for the PB2, PB1, PA, NP, M1 and NS1 genome segments estimated using the sub-sampled alignments. Again, no evidence of reassortment was observed for the HP lineages. Selected HP clusters (C1-C8) are highlighted in red, whilst the HP H5NX (excluding H5N1) sequences within the C8 are highlighted in blue. Bootstrap values are indicated for branches of interest. Clusters are missing in some of the phylogenies is due to the lack of corresponding sequences for all viral genome segments.

**Supplementary Text 5**

### Amino Acid Trait Association Model

#### Model description

We developed a pair of nested evolutionary models, a null and an alternative model, to test for associations between a target amino acid at an alignment site and a target binary trait. The target amino acid being tested, denoted $targetaa$, represents one of the 20 possible amino acids. The target trait, denoted $\mathcal{T}$, represents the trait being tested for an association with the target amino acid. The non-target trait is denoted $\mathcal{N}$. The traits, like the amino acids, are assumed to have evolved in an evolutionary manner.

The null model treats the amino acid evolution at a site and the trait evolution as independent of one another, whereas the alternative model treats the target amino acid and the target binary traits as potentially associated. This potential association is introduced into the alternative model via a dependence parameter $\lambda$.

For a given amino acid site and set of traits we used maximum likelihood estimation to estimate the parameters of both models and to obtain the maximum likelihood values. The maximum likelihood values were used to compare both models using a likelihood ratio test (LRT) and to calculate a p-value. If the LRT rejects the null model ($p < 0.05$) in favour of the alternative model, this suggests that the target amino acid and target trait are associated.

This association can be a positive association: the target amino acid and the target trait tend to co-occur together, or a negative association: the target amino acid and the target trait tend to actively avoid co-occurring together. When the maximum likelihood estimate for the dependence parameter is larger than one, $\hat{\lambda} > 1$, this suggests a positive association, and when $\hat{\lambda} < 1$, this suggests a negative association.

This test is somewhat analogous to a chi-squared test of association, except it accounts for phylogenetic correlations. A chi-squared test will treat each observation of amino acid and trait at the tips of a phylogeny as independent events, when in reality they are produced by an evolutionary process where the underlying number of events leading to those observations may be small. This is commonly referred to as a founder effect (Bhattacharya *et al.*, 2007), and can result in chi-squared associations whose significance is erroneously inflated. Our test avoids this by explicitly modelling the potential dependence between the amino acid and trait evolutionary processes and testing its significance relative to a model that treats them as independent of one another.

Note that the traits, like the amino acids, are assumed to evolve in an evolutionary manner along the tree, and therefore our test is only appropriate where the trait can be described by an evolutionary process. This is the case for the HP and LP traits because they are a direct function of the presence or absence of an insertion, which is generated by an insertion-deletion evolutionary process along the tree. This test would not be appropriate for a trait such as patient survival, which represents a propensity along the tree rather than propagating in a discrete manner. For traits such as these we recommend the test outlined in Bhattacharya *et al.* (2007).

A formal description of the model is given as follows: the joint evolution of amino acids and traits are modelled using a $40 \times 40$ substitution model $Q$, that combines a $20 \times 20$ amino substitution model, $A$, and $2 \times 2$ by trait model, T. The trait model is a two-state continuous-time Markov model akin to Felsenstein's 1981 DNA substitution model (Felsenstein, 1981). The joint model is given as follows:

$$Q_{ij,mn} = \begin{cases} \mu A_{ij}\lambda & \text{if } i \neq j \text{ and } j = targetaa \text{ and } m = n = \mathcal{T} \\ \mu A_{ij}\frac{1}{\lambda} & \text{if } i \neq j \text{ and } j \neq targetaa \text{ and } m = n = \mathcal{T} \\ \mu A_{ij} & \text{if } i \neq j \text{ and } m = n = \mathcal{N} \\ \pi_{\mathcal{T}}\tau\lambda & \text{if } i = j = targetaa \text{ and } m = \mathcal{N} \text{ and } n = \mathcal{T} \\ \pi_{\mathcal{T}}\tau\frac{1}{\lambda} & \text{if } i = j \text{ and } j \neq target\text{aa and } m = \mathcal{N} \text{ and } n = \mathcal{T} \\ \pi_{\mathcal{N}}\tau & \text{if } i = j \text{ and } m = \mathcal{T} \text{ and } n = \mathcal{N} \\ 0 & \text{otherwise} \end{cases} \tag{1}$$

Where $i$ and $j$ represent the initial and end amino acid states, respectively, and $m$ and $n$ represent the start and end trait states, respectively. This formulation is motivated by the RNA base-pairing model of Muse (1995).

$\mu$ is a site-specific amino acid substitution rate, and $\tau$ is the trait substitution rate. $A$ is a rate matrix given by the LG2008 substitution model. $\pi^A$ is a vector of 20 amino acid frequencies as specified by the LG2008 model, and $\pi_{\mathcal{T}}$ and $\pi_{\mathcal{N}}$ represents the frequencies of the non-target ($\mathcal{T}$) and target traits ($\mathcal{N}$) states, respectively.

The equilibrium frequencies, $\pi$, of the alternative model are given by four separate cases corresponding to the two possible values for traits ($\mathcal{T}$ or $\mathcal{N}$), and whether the amino acid ($aa$) matches the target amino acid ($targetaa$) or not:

$$\pi_{aa=targetaa,trait=\mathcal{T}} = k^{-1}\pi^A_{aa}\pi_{\mathcal{T}}\lambda \tag{2}$$

$$\pi_{aa\neq targetaa,trait=\mathcal{T}} = k^{-1}\pi^A_{aa}\pi_{\mathcal{T}}\frac{1}{\lambda} \tag{3}$$

$$\pi_{aa=targetaa,trait=\mathcal{N}} = k^{-1}\pi^{A}_{aa}\pi_{\mathcal{N}} \quad (4)$$

$$\pi_{aa\neq targetaa,trait=\mathcal{N}} = k^{-1}\pi^{A}_{aa}\pi_{\mathcal{N}} \quad (5)$$

Where $\kappa = (\lambda + \frac{1}{\lambda})\pi_{\mathcal{T}} + 2\pi_{\mathcal{N}}$ is a normalising constant.

These equilibrium frequencies provide a intuitive way of understanding the influence of the association parameter $\lambda$. It is possible to get a sense of the expected frequencies of particular amino acid and trait associations for given values of $\lambda$. Furthermore, they can be used to predict, for a single sequence, the posterior probability of a trait given the amino $aa$ at the target site:

$$p(trait = \mathcal{T}|aa = targetaa) = \frac{\pi_{\mathcal{T}}\lambda}{\pi_{\mathcal{N}} + \pi_{\mathcal{T}}\lambda} \quad (6)$$

$$p(trait = \mathcal{T}|aa \neq targetaa) = \frac{\pi_{\mathcal{T}}}{\pi_{\mathcal{N}}\lambda + \pi_{\mathcal{T}}} \quad (7)$$

Also note that the model is time-reversible, and therefore an unrooted tree can be used if the equilibrium probabilities are taken to be the initial probabilities at any rooting of the tree (Felsenstein, 1981).

#### Simulations

Table 1: Summary of benchmarks results

| **Simulated association strength** | **Simulated rate of trait evolution** | **Recall** | **Precision** |
|---|---|---|---|
| Weak (2.0) | 2.0 | 0.23 | 0.94 |
| Intermediate (4.0) | 2.0 | 0.35 | 0.96 |
| Strong (8.0) | 2.0 | 0.50 | 0.93 |
| Weak (2.0) | 4.0 | 0.26 | 0.92 |
| Intermediate (4.0) | 4.0 | 0.48 | 0.92 |
| Strong (8.0) | 4.0 | 0.64 | 0.96 |
| Weak (2.0) | 7.5 | 0.28 | 0.97 |
| Intermediate (4.0) | 7.5 | 0.62 | 0.98 |
| Strong (8.0) | 7.5 | 0.73 | 0.95 |

We simulated amino acid alignments using the empirical H7 AIV maximum likelihood tree and its corresponding HP and LP trait values. Each alignment consisted of 500 amino acids, with the first twenty sites of each alignment simulated as being associated with the traits, each having a different one of the 20 canonical amino acids as the target amino

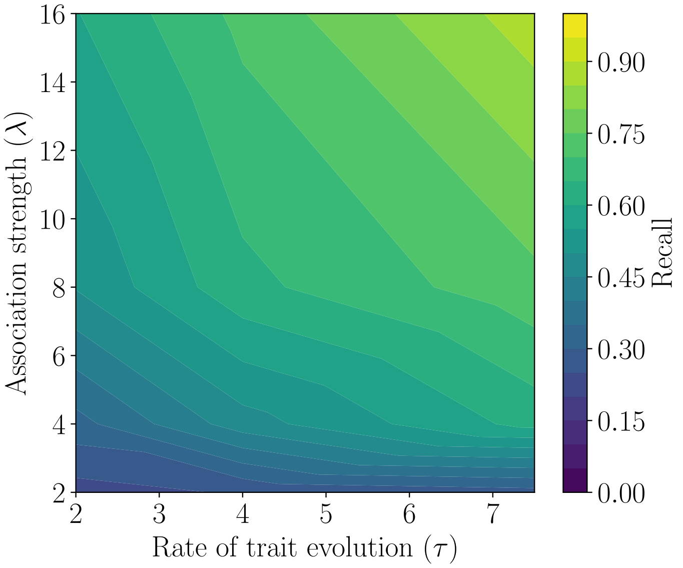


Figure 1: Contour plot of recall (blue-green-yellow colour gradient) as a function of simulated rate of trait evolution (x-axis) and simulated association strength (y-axis).

acid. The remaining 480 amino acid sites were simulated under the LG2008 model and were therefore treated as being independent of the traits. Three different degrees of association were simulated: weak, intermediate, and high, combined with three different rates of trait evolution (2.0, 4.0, and 7.0) - the inferred rate of trait evolution in the H7 ML tree was $\sim$4.0 and so this was selected as an intermediate value.

To account for the potential error introduced during tree inference, an ML tree was inferred using FastTree (Price *et al.*, 2010) for each of the simulated alignments. Potential associations were then estimated using our model on the first 40 sites of each alignment. The first twenty sites were used to measure the number of true positive and false negative detections, whereas the remaining twenty sites were simulated as independent of the traits and were used to measure the number of true negative and false positive detections. The recall and precision were calculated for each simulation using the number of true positives (TN), false-positives (FP), and false-negatives (FN). Recall and precision are defined as follows:

$$\text{Recall} = \frac{\text{TP}}{\text{TP} + \text{FN}} \tag{8}$$

$$\text{Precision} = \frac{\text{TP}}{\text{TP} + \text{FP}} \tag{9}$$

The results in Table 1 indicate that our model has a false-discovery rate (FDR= $100\% \times [1.0-$

Precision]) of $\sim 5\%$ across all test conditions which is consistent with our p-value significance threshold of 0.05. The recall of the model increases with the simulated association strength as expected, with stronger associations being more easily detected. Likewise, the recall of the model increases with a higher rate of trait evolution (Table 1 and Figure 1), which is also expected given that higher rates of trait evolution imply a greater number of trait events along the tree and therefore more the power in being able to detect an association.

##### Software availability

Julia source code (compatible with Windows and Linux) is available at:
https://github.com/michaelgoldendev/trait-evolution

#### References

Bhattacharya, T., Daniels, M., Heckerman, D., Foley, B., Frahm, N., Kadie, C., Carlson, J., Yusim, K., McMahon, B., Gaschen, B., *et al.* 2007. Founder effects in the assessment of hiv polymorphisms and hla allele associations. *Science*, 315(5818): 1583–1586.

Felsenstein, J. 1981. Evolutionary trees from DNA sequences: a maximum likelihood approach. *Journal of molecular evolution*, 17(6): 368–376.

Muse, S. V. 1995. Evolutionary analyses of DNA sequences subject to constraints of secondary structure. *Genetics*, 139(3): 1429–1439.

Price, M. N., Dehal, P. S., and Arkin, A. P. 2010. FastTree 2–approximately maximum-likelihood trees for large alignments. *PloS one*, 5(3): e9490.
